# Investigation of the anti-TB potential of selected alkaloid constituents using molecular docking approach

**DOI:** 10.1101/2020.04.28.067090

**Authors:** Mohammad Kawsar Sharif Siam, Mohammad Umer Sharif Shohan, Zaira Zafroon

## Abstract

*Mycobacterium tuberculosis*, the leading bacterial killer disease worldwide, causes Human tuberculosis (TB). Due to the growing problem of drug resistant *Mycobacterium tuberculosis* strains, new anti-TB drugs are urgently needed. Natural sources such as plant extracts have long played an important role in tuberculosis management and can be used as a template to design new drugs. A wide screening of natural sources is time consuming but the process can be significantly sped up using molecular docking. In this study, we used a molecular docking approach to investigate the interactions between selected natural constituents and three proteins MtPanK, MtDprE1 and MtKasA involved in the physiological functions of Mycobacterium tuberculosis which are necessary for the bacteria to survive and cause disease. The molecular docking score, a score that accounts for the binding affinity between a ligand and a target protein, for each protein was calculated against 150 chemical constituents of different classes to estimate the binding free energy. The docking scores represent the binding free energy. The best docking scores indicates the highest ligand protein binding which is indicated by the lowest energy value. Among the natural constituents, Shermilamine B showed a docking score of - 8.5kcal/mol, Brachystamide B showed a docking score of −8.6 kcal/mol with MtPanK, Monoamphilectine A showed a score of −9.8kcal/mol with MtDprE1.These three compounds showed docking scores which were superior to the control inhibitors and represent the opportunity of in vitro biological evaluation and anti-TB drug design. Consequently, all these compounds belonged to the alkaloid class. Specific interactions were studied to further understand the nature of intermolecular bonds between the most active ligands and the protein binding site residues which proved that among the constituents monoamphilectine A and Shermilamine B show more promise as Anti-TB drugs. Furthermore, the ADMET properties of these compounds or ligands showed that they have no corrosive or carcinogenic parameters.

**Graphical Abstract:** 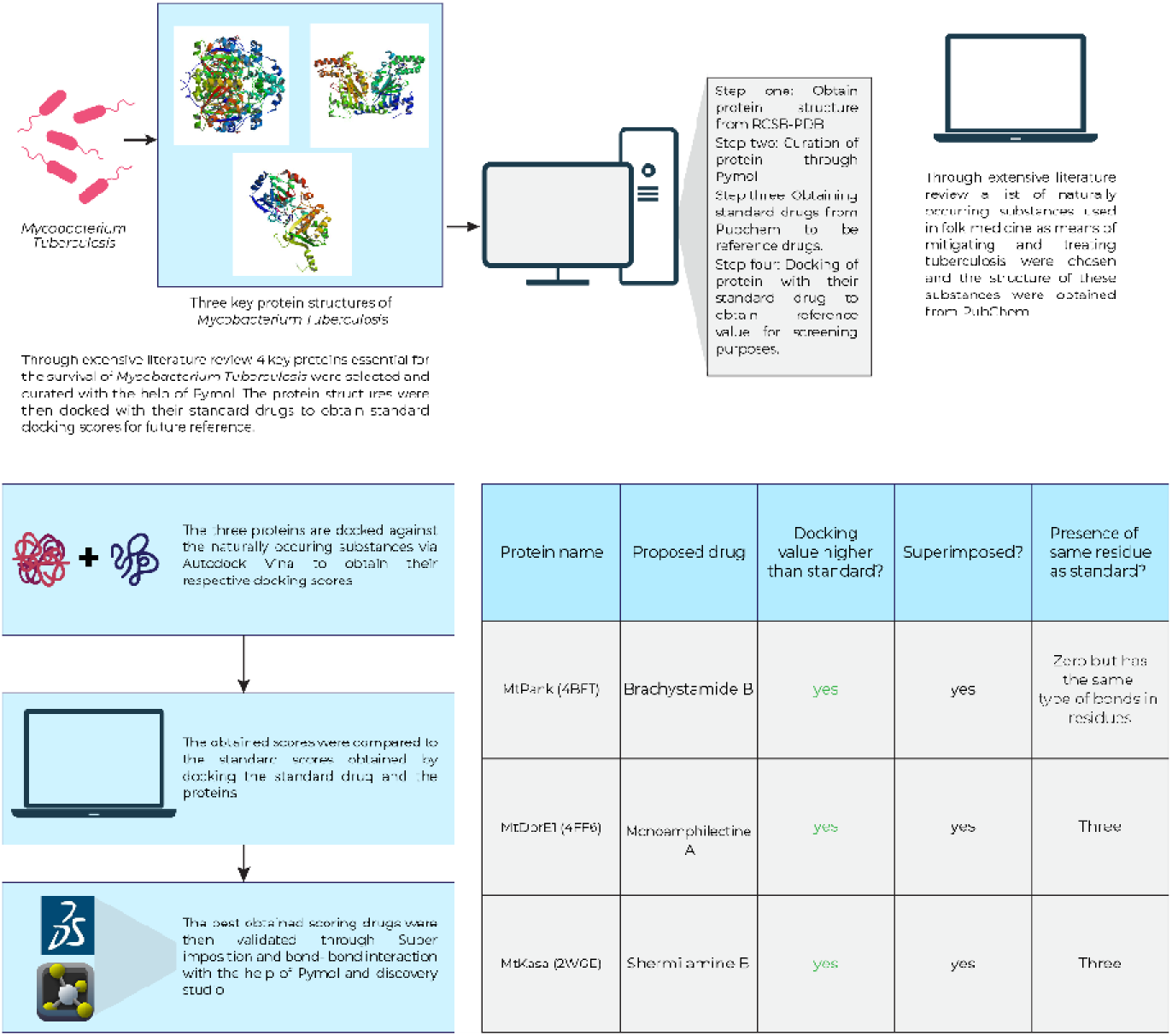

## 1. Introduction

Tuberculosis (TB), primarily caused by *Mycobacterium tuberculosis*, is a major disease that affects millions of people each year with high mortality rates. The ‘End-Tb Strategy’ goal of the World Health Organization and the Sustainable Development Goals (SDGs) of the United Nations (Goal 3; goal 3) lay out the roadmap for achieving a global goal to end the Tb epidemic by 2030. The lack of treatment options accompanied by the current emergence of MTB’s multi-drug resistant (MDR) and extreme drug resistance (XDR) strains [1] [2] remains a barrier in solving this problem.[3][4] There are a low number of drugs available for the treatment of Tb (MDR / XDR) and there are several reasons for the lack of new drugs, including the lack of funding for such neglected diseases in Pharmaceutical Research & Development. Drug development’s prohibitive cost has been caused by poor target selection and as a result, 87 percent of late-stage failures can be avoided as they show poor effectiveness and negative effects [5]. The lessons learned from current efforts to generate TB hits and newly validated drug targets for TB can now be applied to generate new TB drugs. Using currently underexploited chemicals sources and lead-optimization methods can improve the efficiency of drug development process.

Historical findings and reports show that plants were used in earlier era for medical purposes [6]. Mdluli discusses probable leads for the development of antimicrobial drugs in his review. For anticancer and antimicrobials drugs in particular, natural products have been the go to source of dug compounds [7][8][9]. A number of researchers have highlighted the urge to discover and develop new antitubercular drugs to reduce the burden of deadly disease, also known as “Captain of Death,”(B. B. Mishra & Tiwari, 2011;Zumla, Nahid, & Cole, 2013). [12]. Various plant extracts were commonly used in traditional medicine for the treatment of tuberculosis and there is a renewed interest in plants as a source of new drug development [13].There is research that proves alkaloids have wide therapeutic applications and can be an ideal target for drug development as researches are being done to develop new drugs from natural sources for tuberculosis [14]. Extracts from Diplosoma and sea squirts lissoclinum showed the presence of an alkaloid called Shermilamine B and inhibits the growth of tuberculosis at the MIC value of mM [15]. An alkyl amide named brachystamide B, has been extracted from *Piper Sarmentosum* and has showed moderate inhibitory activity against M. Tuberculosis at the MIC value of 50 mg/ mL[16]. Monoamphilectine A (diterpenoid b-lactam alkaloid) isolated from the Hymenia cidon marine sponge, at MIC value of 15.3 mg / mL showed potent antimycobacterial properties [17].

Molecular docking can be described as an essential part in the process of computer aided drug design. It is a widely used protocol which is generally used to evaluate the binding affinity of a ligand to its target receptor[18][19] In case of therapeutics and sensor design molecular recognition plays a key role[20]. The role of molecular recognition is particularly important in case of biological systems where they are known to play a role in receptor ligand complex formation[21]. A molecular receptor’s selective binding capability to a ligand with a high affinity requires molecular recognition by non-covalent bonds such as hydrogen bonds, ionic bonds and van der Waals attractions along with hydrophobic interactions[22][23]. Identifying and quantifying these non-covalent interactions which are weak in nature, is essential for structural ligand design[24]. Hydrogen bond in particular is an important factor related to molecular recognition, and binding energy is at the core of all molecular recognition process as well as all processes related to catalysis of enzymes[25]. Intermolecular hydrogen bonding plays a significant role in the field of molecular and supramolecular chemistry, and has critical effects in solvent-solute interaction[26].

Several proteins that play key physiological functions in *M*. *tuberculosis* have been identified as novel attractive molecular targets for anti-TB drug development. The mycobacterial β-ketoacyl ACP synthase known as MtKasA plays a key role in the FAS-II system. KasA is essential for mycobacteria, cell lysis is induced by conditional depletion of KasA [27] and hybridization of the transporon site has shown that KasA plays a key role in survival of the pathogens [28]. Several bacteria have two PanK genes coding for various enzyme types. The genome of *M. Tuberculosis* contains both coaA and coaX genes, which code for type I and type III PanK. However, coaA has been shown to be the only PanK gene essential for in vitro and in vivo bacterial growth [29]. Arabinan is a fundamental part of the pathogen’s cell wall. Decaprenylphosphoryl arabinose, the single donor of arabinosyl residues for the buildup of arabinans is catalyzed by a unique epimerization reaction by the Mycobacterial enzyme DprE1 and DprE2 [30].Thus, these three proteins present novel drug development opportunities.

In this study, we used a molecular docking approach to target the three druggable proteins e.g. Mycobacterium tuberculosis pantothenate kinase (MtPanK, type 1)[31], Mycobacterium tuberculosis decaprenylphosphoryl-β-D-ribose 2′-epimerase 1 (MtDprE1)[32], and β-ketoacyl acyl carrier protein synthase I (MtKasA)[33][34] with alkaloid compounds using Auto dock Vina. Auto dock Vina is largely used to screen potential of small ligands in inhibiting the activities of the proteins including docking of alkaloid with *mycobacterium tuberculosis* proteins. [35]–[40]. Furthermore, all the interactions were further studied to understand the bond nature of these molecules with the protein to elucidate their mechanism of action when compared to the chosen standard drugs used to inhibit them. Additionally, bond types were studied to ensure that they have biological importance. Moreover, absorption, metabolism and toxicity analysis of these small ligands were performed to assess their pharmacokinetic properties which indicated that these compounds are non-carcinogenic and possess no harm to humans.[41]. There have been no reports of docking of alkaloids towards MtPanK, MtDprE1, and MtKasA to our knowledge.

## 2. Materials and Methods

### 2.1 Ligand selection

The ligands selected for this study were 150 well characterized phytochemicals previously used in traditional medicine [8], [14], [42], [43]. All chemical structures were retrieved from the PubChem compound database (NCBI) (http://www.pubchem.ncbi.nlm.nih.gov).

### 2.2 Ligand and protein preparation

Each ligand structure was obtained from PubChem Database and open babel was used to turn the. sdf file to. pdb file[39].After which the ligand was opened in Auto dock Vina and the existing rotatable bonds present on the ligands were treated as non-rotatable. This allowed rigid docking to be performed and minimize the chance of standard errors[44]. Gasteiger charges were calculated and before docking partial charges were attached to the ligands [45]. The crystal structures of MtPanK type 1 (PDB ID: 4BFT), MtDprE1 (PDB ID: 4FF6), and MtKasA (PDB ID: 2WGE) were retrieved from the RCSB Protein Data Bank (PDB) database (http://www.pdb.org). The structures of the ligand inhibitors 2-chloro-N-[1-(5-{[2-(4-fluorophenoxy)ethyl] sulfanyl}-4-methyl-4h-1,2,4-triazol-3-Yl) ethyl]benzamide (ZVT) which is the standard inhibitor for MtPanK, 3-(hydroxyamino)-N-[(1r)-1-phenylethyl]-5-(trifluoromethyl)benzamide (OT4) which is the standard inhibitor for MtDprE1, and thiolactomycin (TLM) which is the standard inhibitor for MtKasA were retrieved from their corresponding PDB entries. Each protein was used as a rigid structure and all water molecules and hetero-atoms were removed using Pymol [46].

### 2.3 Identification of binding site residues

Previous studies were used to identify the nature and the role of the binding site residues for MtPanK, MtDprE1 and MtKasA. Specific amino acids involved in ligand/protein interactions were observed by visualizing the control inhibitor and protein interactions in BIOVIA Discovery Studio Visualizer v.4.5.[47]

### 2.4 Grid box preparation and docking

The open source chemical toolbox Open Babel v. 2.3.283 was used to perform all file conversions needed prior to docking[39]. The parameters for the grid box was set in a way to form a sufficiently big cavity room within the binding site of each protein to accommodate each compound and the cavity parameters were determined using Auto dock Tools v. 1.5.6rc384. Molecular docking calculations were conducted using Auto dock Vina v. 1.1.281 for all compounds with each of the proteins. To dock the compounds against the four proteins, the center grid box was positioned at the center of the protein structure and was expanded in x, y and z directions until the grid box fully covered the protein structure. The Auto dock Vina protocol was used to dock the compounds in respect to the proteins and to predict binding affinities between the drug receptor complex[35]. To validate precision of docking results, all inhibitory co-crystallized ligands have been re-docked into the respective protein structures that they were removed from. The algorithms of Auto dock vina searched and ranked different orientations of the ligands based on the energy scores. In the co-crystallized protein-ligand complex, our docking protocol was able to create a comparable docking pose for each control ligand to its biological confirmation. We further inspected all binding poses for a given ligand visually and only poses with the lowest RMSD value (simply root-mean-square deviation) (threshold < 1.00 Å) were deemed to achieve a greater docking precision. During the docking phase, the Lamarckian Genetic Algorithm was used to investigate the best conformational space for each 150-population ligand. The maximum numbers of generation was set at 27,000 and evaluation was set at 2,500,000.rest of the parameters were set as default. We decided to use a guided docking strategy to improve docking efficiency [38] and ranked the conformations based on their predicted binding affinities(Table S1). The lower the binding free energy the higher the predicted ligand-protein affinity. The Auto dock Vina docking scores of these selected alkaloid constituents that ranked higher than a control inhibitor was further studied in Pymol. Specific intermolecular interactions with the targets (Table S7-S9 & Figs 1(a, b, c) were further visualized using BIOVIA Discovery Studio Visualizer v.4.5 (Accelrys). The ADMET properties were checked using ADMETSAR 2.0 to ensure that the alkaloids were not carcinogenic[48].

**Figure 1:**
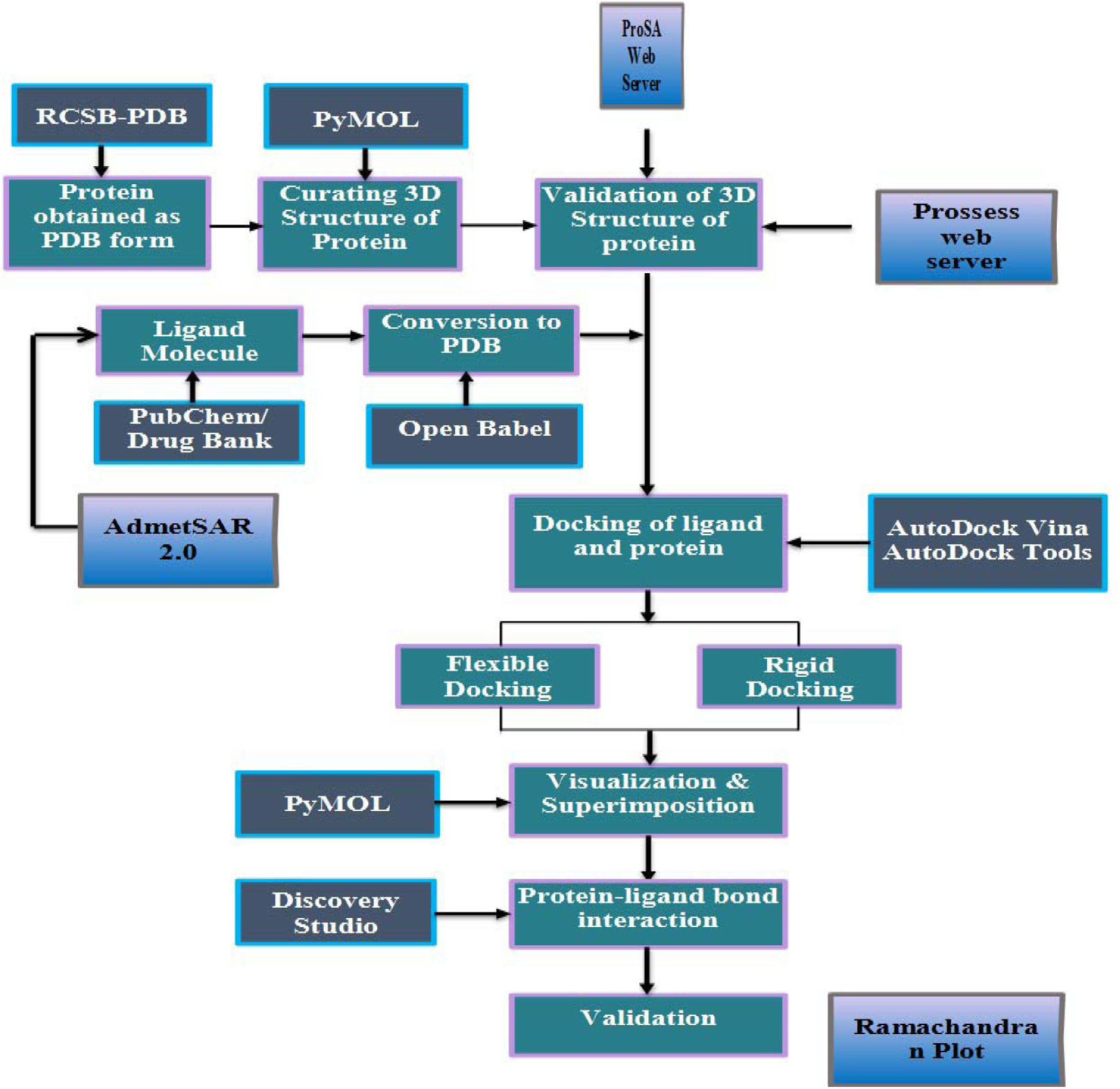
Methodology

The entire process has been illustrated below:

## 3. Results

150 chemical constituents of four different classes were selected based on their traditional use for Tuberculosis in ancient medicine. Molecules earlier identified as target enzyme inhibitors in the literature, were used as controls. They were chosen as controls due to the fact that the nature and the role of binding site residues were known from the available protein ligand complex. To reaffirm the docking parameters all, control inhibitors were removed from its co-crystalized complex forms with the help of Pymol. After which the ligands were re-docked with the help of Auto dock Vina against the chosen enzymes. A docking score for alkaloids and other classes of small compounds was calculated with Auto dock vina to estimate the binding free energy towards MtKasA, MtPanK, and MtDprE1. (Table 1) To choose molecules with minimum energy values that ranked higher than the control inhibitors selected alkaloids were docked against the target proteins and the docking scores obtained for compounds within each phytochemical class were compared to the scores of the control inhibitors for each target. It was seen that alkaloids performed better than the other phytochemical classes. We observed that three alkaloid constituents exhibited scores that ranked better than the controls against MtPanK MtKasA and MtDprE1. Shermilamine B and brachystamide B showed strong docking scores towards MtKasA (−8.5 and −8.4 kcal/mol, respectively) superior to the control inhibitor thiolactomycin (−7.9 kcal/mol). Shermilamine B, brachystamide B and monoamphilectine A also showed strong predicted binding towards MtDprE1 (−9.7, −9.7 and-9.8 kcal/mol) compared with the control inhibitor 0T4 (−9.2 kcal/mol). Brachystamide B and monoamphilectine A also showed strong predicted binding towards MtPanK (−8.6, −7.5 kcal/mol) which is higher when compared to the control inhibitor ZVT (−7.3 kcal/mol). Using Pymol[46], [49], superimposition results were observed for control inhibitor and proposed constituents.

**Table 1:**
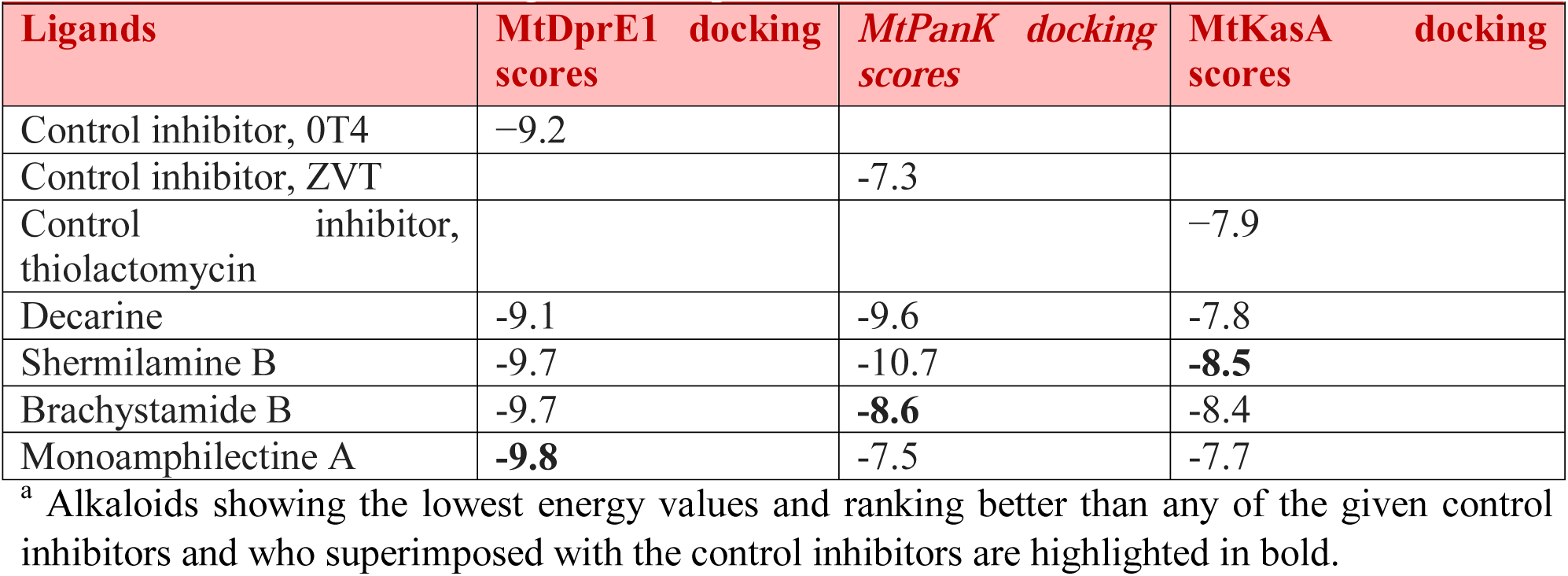
Predicted binding affinity (docking scores in kcal/mol)) of selected alkaloids and re-docked control inhibitors against MtDprE1, MtKasAa.and MtPanK^a^

|  |  |  |  |
| --- | --- | --- | --- |
| Ligands | MtDprE1 docking scores | MtPanK docking scores | MtKasA docking scores |
| Control inhibitor, OT4 | −9.2 |  |  |
| Control inhibitor, ZVT |  | -7.3 |  |
| Control inhibitor, thiolactomycin |  |  | −7.9 |
| Decarine | -9.1 | -9.6 | -7.8 |
| Shermilamine B | -9.7 | -10.7 | **-8.5** |
| Brachystamide B | -9.7 | **-8.6** | -8.4 |
| Monoamphilectine A | **-9.8** | -7.5 | -7.7 |

<sup>a</sup> Alkaloids showing the lowest energy values and ranking better than any of the given control inhibitors and who superimposed with the control inhibitors are highlighted in bold.

The superimposition results are presented in the supplementary Table S1. Specific interactions were evaluated to study the nature of the intermolecular bonds formed between the chosen compounds and the binding site residues of the three proteins that were studied. (Table S2-S4)

The binding poses obtained for the best binding ligands Shermilamine B, brachystamide B and monoamphilectine A were visually inspected and are depicted in Figure 1 (a, b, c) respectively. Furthermore, the Adsorption, distribution, metabolism, excretion, toxicity of the alkaloid compounds was measured to ensure that the drugs did not have any carcinogenic property and did not cause acute toxicity.[48] Table S5 – S7

**Figure 2:**
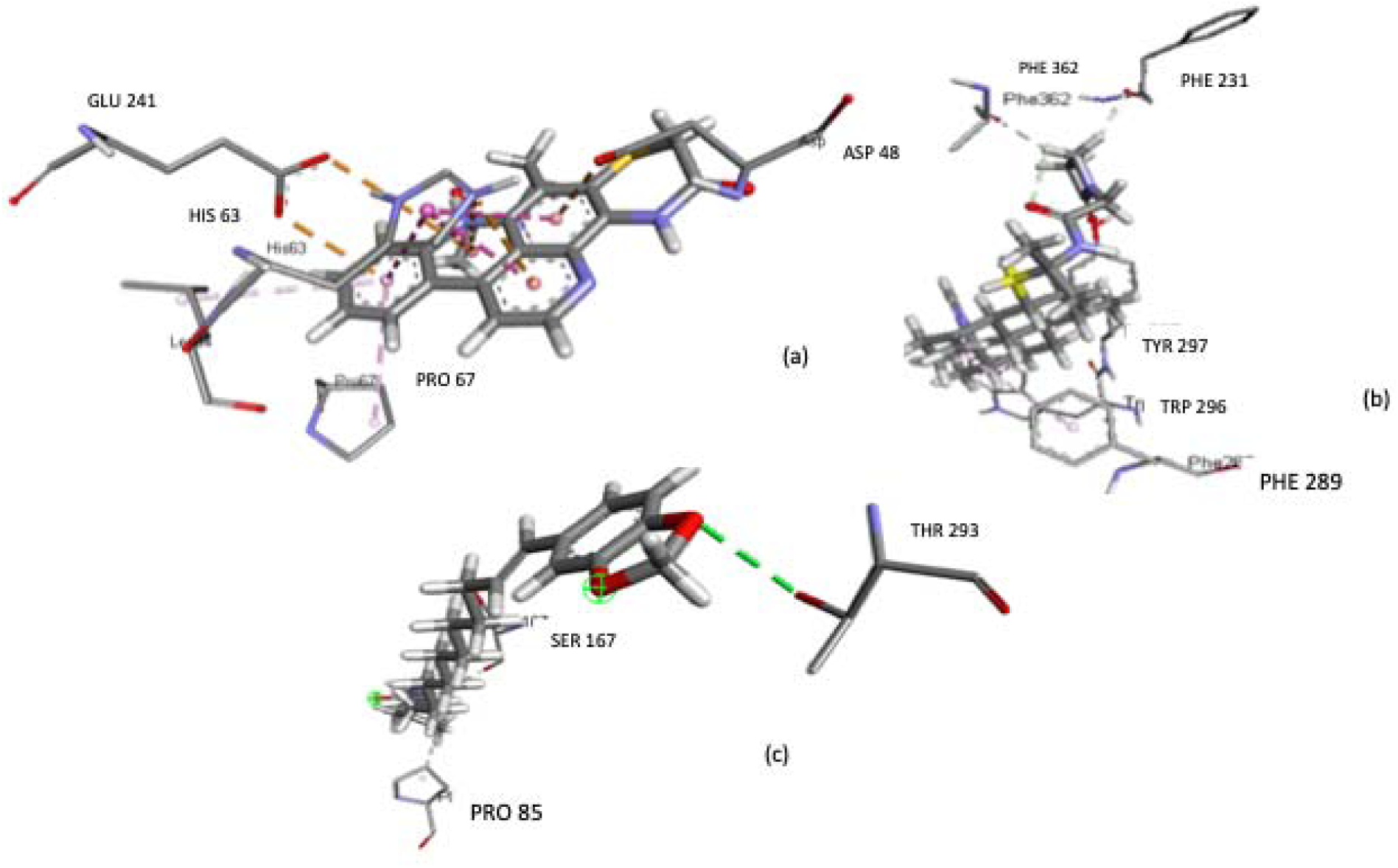
2D plot of interactions between *(a)* MtKasA and Shermilamine B (b) MtDprE1 and Monoamphilectine A (c) MtPanK and brachystamide B generated by BIOVIA Discovery Studio visualizer.

## 4. Discussion

MtKasA, MtPanK and MtDprE1 are proteins, which are essential for the growth, pathogenicity and survival of mycobacterium tuberculosis. As a major controller of the developmental processes in tuberculosis, these proteins can be considered as a therapeutic target. These proteins are also absent in the mammalian species thus these proteins are highly selective and can be used as target for inhibition of tuberculosis.[28]–[31], [50], [51]. The cell wall of mycobacteria is a complicated structure that is made up of lipoarabinomannan, peptidoglycan, mycolic acids and arabinogalactan. Thus, two proteins that play a key role in the synthesis of the bacterial cell wall was used in the study namely β-ketoacyl acyl protein synthase I (MtKasA) and decaprenylphosphoryl-β-D-ribose 2′-epimerase 1 (MtDprE1)[32], [34]. Type I pantothenate kinase of Tuberculosis (MtPanK) is a key enzyme which is involved in the catalyzation of cofactor Coenzyme A (CoA) biosynthesis through converting pantothenateto 4′-phosphopantothenate [31].

After doing an extensive review on proteins associated with tuberculosis, tuberculosis and computational techniques this study was designed to find out some potential drugs to inhibit key proteins associated with the survival of *M. Tuberculosis.* More than 150 random chemical constituents derived from natural sources of different classes were selected for this study to be evaluated to be potential anti TB drugs. Throughout the study, several in silico techniques were used. The selected target proteins and their known established inhibitors were docked against each other to find the standard docking score which was used as control to evaluate the potential of the selected natural compounds.

After docking all the compounds, the binding affinities of alkaloids after rigid docking using autodock vina were found between −8.5 kcal/mol to −9.8 kcal/mol that represents a strong binding affinity. The binding affinities of the established inhibitors were also calculated using same method. The results of their binding affinities were higher for three of the target proteins (MtKasA, MtPanK, MtDprE1) compared to their control or standard inhibitor. The choice of drug to target the MtKasA protein is Shermilamine B. Shermilamine B showed a docking score of −8.5kcal/mol, which was higher than the standard TLM score. The choice of drug to target the MtPanK protein is Brachystamide B. Brachystamide B showed a docking score of –8.6 kcal/mol that was higher than the standard ZVT score. The choice of drug to target the MtDprE1 protein is monoamphilectine A. Monoamphilectine A showed a score of −9.8kcal/mol which is higher than the standard score of 0T4.Therefore, it is clear that the chosen drugs have stronger affinity towards the target proteins and can be used as potential leads. All the lead compounds have previously shown activities against *M. Tuberculosis.*

After visualizing the protein and alkaloid and protein and standard drug binding using PyMOL, it was found out that the alkaloid constituents and established inhibitors bind within a same pocket in the target proteins.

In order to know the protein-ligand interactions of all the ligands with proteins, Discovery Studio Visualizer was used. Upon the observation of the results, it was observed that three of the residues GLU 241, GLU 241: OE1, GLU 242: OE2 and HIS 63 residues are common interacting residues of the control inhibitor (TLM) and proposed alkaloid (Shermilamine B) when bound to MtKasA. It was also observed that two of the residues TYR 297 and PHE 362 residues are common interacting residues of the control inhibitor (OT4) and proposed alkaloid (monoamphilectine A) when bound to MtDprE1.Lastly, it was observed that there are no same interacting residues between the control inhibitor and proposed alkaloid brachystamide B. It is also known that hydrogen bonds are important in case of binding of a drug or ligand to a receptor [52], [53] The pi cation bonds plays an important role in molecular recognition and chemical and biological catalysis[54].On top of this, using ADMETSAR 2.0, it was seen that none of these alkaloids are carcinogenic[48].

Based on interaction studies it can be seen that strong bonds such as hydrogen bonds and pi cation and pi anion bonds featured significantly in the results proving that these ligands can be potential drugs in the future namely MtDprE1 can be targeted using monoamphilectine A, MtKasA can be targeted using Shermilamine B, and MtPanK can be targeted using brachystamide B.

## 5. Conclusion

It is an established fact that tuberculosis has become a pandemic disease and new drugs are an urgent need. In the study, we used a molecular docking approach to identify potential lead compounds for the treatment of tuberculosis and were able to confirm three alkaloids as potential candidates for development of new multidrug resistant compounds. As these alkaloids show great promise in vivo studies and generate multiple hit they can also be further structurally optimized for the design of new anti-TB drugs. Further, in vitro studies are necessary for this.

## List of Abbreviations

ADMET: Absorption, Distribution, Metabolism, Excretion and Toxicity
TB: Tuberculosis
MTB: Mycobacterium tuberculosis
MDR: Multi-drug resistant
XDR: Extreme drug resistance
MtPanK: Mycobacterium tuberculosis pantothenate kinase
MtDprE1: Mycobacterium tuberculosis decaprenylphosphoryl-β-D-ribose 2′-epimerase 1
MtKasA: Mycobacterium tuberculosis β-ketoacyl acyl carrier protein synthase I
ZVT: 2-chloro-N-[1-(5-{[2-(4-fluorophenoxy)ethyl] sulfanyl}-4-methyl-4h-1,2,4-triazol-3-Yl) ethyl]benzamide
OT4: 3-(hydroxyamino)-N-[(1r)-1-phenylethyl]-5-(trifluoromethyl)benzamide
TLM: Thiolactomycin

## 6. Acknowledgment

This research received no external funding.

## 7. Declaration of Competing Interest

The author(s) declared no potential conflicts of interest with respect to the research, authorship, and/or publication of this article.

## 8. Author Contributions

Mohammad Kawsar Sharif Siam conceived, designed and supervised the experiments, performed the experiments, analyzed the data, contributed materials/analysis tools, prepared figures and/or tables, authored or reviewed drafts of the paper, approved the final draft.

Mohammad Umer Sharif Shohan conceived and designed the experiments, performed the experiments, analyzed the data, authored or reviewed drafts of the paper, approved the final draft.

Zaira Zafroon analyzed the data, authored or reviewed drafts of the paper, approved the final draft.

## Notes

### Competing Interest Statement

The authors have declared no competing interest.

