## Supplementary material for "Investigation of the anti-TB potential of selected alkaloid constituents using molecular docking approach": Table S1, Table S2 - S4, Table S5 - S7

| **Table S1: Superimposition results for alkaloids and control inhibitors** | | |
| --- | --- | --- |
| **Superimposition results** | **Drugs** | **Docking Results**  **(Binding Affinity (kcal/mol)** |
| 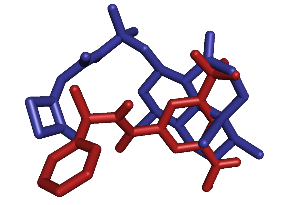 | 0T4( control inhibitor) | -9.2 |
|  | Monoamphilectine A | -9.8 |
|  | Remarks: PyMol was used to view superimposition results of Monoamphilcetine A and control inhibitorOT4. The target protein was MtDprE1(4FF6) | |

**Supplementary tables:**

**Investigation of the anti-TB potential of selected alkaloid constituents using molecular docking approach**

Mohammad Kawsar Sharif Siam^1*^, Mohammad Umer Sharif Shohan^2^, Zaira Zafroon^1^

^1^ Department of Pharmacy, BRAC University, Dhaka, Bangladesh

^2^ Department of Biochemistry and Molecular Biology, University of Dhaka, Dhaka, Bangladesh

Table S1: Superimposition results for alkaloids and control inhibitors

| **Superimposition results** | **Drugs** | **Docking Results**  **(Binding Affinity (kcal/mol)** |
| --- | --- | --- |
| 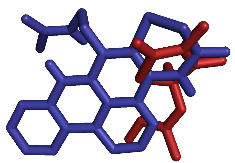 | TLM( control inhibitor) | -7.9 |
|  | Shermilamine B | -8.5 |
|  | Remarks: PyMol was used to view superimposition results of shermilamine B and control inhibitor TLM. The target protein was MtKasA(2WGE) | |

| **Superimposition results** | **Drugs** | **Docking Results**  **(Binding Affinity (kcal/mol)** |
| --- | --- | --- |
| 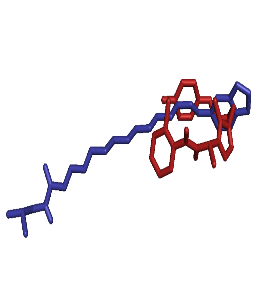 | ZVT( control inhibitor) | -7.3 |
|  | BRACHYSTAMIDE B | -8.6 |
|  | Remarks: PyMol was used to view superimposition results of brcahystamide B and control inhibitor ZVT. The target protein was MtPanK(4BFT) | |

Table S2: Docking against MtDprE1 (PDB ID- 4ff6) and their Discovery studio visualization results

| **Table S2: Docking against MtDprE1 (PDB ID- 4ff6) and their Discovery studio visualization results** | | | | | | | |
| --- | --- | --- | --- | --- | --- | --- | --- |
| **Compound Name** | **Docking against MtDprE1 (PDB ID- 4ff6)** | | | | | | |
|  | Binding energy  (kcal/mol) | H-bond | | Hydrophobic Interaction | | Electrostatic Interaction | |
|  |  | (Amino acid-ligand atom) | Distance  (Å) | (Amino acid-ligand atom) | Distance  (Å) | (Amino acid-ligand atom) | Distance  (Å) |
| **OT4**  (Standard Drug) | -9.2 | TYR297: | 2.42752 | THR288 | 3.88147 | ASP232 | 4.89466 |
|  |  | PHE362 | 3.09613 | ILE234 | 4.71427 |  |  |
| **Monoamphilectine A** (Proposed Alkaloid) | -9.8 | PH | 2.20582 | PHE289 | 4.79301 | - | - |
|  |  | PHE362 | 2.31394 | TRP296 | 5.37054 |  |  |
|  |  | H37 | 2.93292 | TRP296 | 4.32284 |  |  |
|  |  |  |  | TYR297 | 4.69661 |  |  |
|  |  |  |  | TYR297 | 4.02288 |  |  |

**Table S3: Docking against MtPanK (PDB ID- 4BFT) and their Discovery studio visualization results**

| **Table S3: Docking against MtPanK (PDB ID- 4BFT) and their Discovery studio visualization results** | | | | | | | |
| --- | --- | --- | --- | --- | --- | --- | --- |
| **Drugs Name** | **Docking against MtPanK (PDB ID- 4BFT)** | | | | | | |
|  | Binding energy  (kcal/mol) | H-bond | | Hydrophobic Interaction | | Electrostatic Interaction | |
|  |  | (Amino acid-ligand atom) | Distance  (Å) | (Amino acid-ligand atom) | Distance  (Å) | (Amino acid-ligand atom) | Distance  (Å) |
| **ZVT** (Standard Drug) | -7.3 | ARG67 | 3.36209 | ARG310 | 5.25027 | - - | |
|  |  | ASP219 | 2.78571 |  |  |  |  |
| **Brachystamide B** (Proposed Drug) | -8.6 | SER167 | 3.00463 | PRO85 | 3.91669 | - | - |
|  |  | THR293 | 2.01872 |  |  |  |  |

**Table S4: Docking against MtKasA (PDB ID- 2WGE) and their Discovery studio visualization results**

| **Table S4: Docking against MtKasA (PDB ID- 2WGE) and their Discovery studio visualization results** | | | | | | | | |
| --- | --- | --- | --- | --- | --- | --- | --- | --- |
| **Compound Name** | **Docking against** | | | | | | | |
|  | Binding energy  (kcal/mol) | H-bond | | Hydrophobic Interaction | | Electrostatic Interaction | | |
|  |  | (Amino acid-ligand atom) | Distance  (Å) | (Amino acid-ligand atom) | Distance  (Å) | (Amino acid-ligand atom) | Distance  (Å) | |
| **Thiolactomycin**  (Standard Drug) | -7.9 | GLU 241 | 2.79986 | PRO 201 | 4.28569 | ASP232 | 4.89466 | |
|  |  | GLU 241 | 2.93651 | PRO 201 | 4.283 |  |  |  |
|  |  |  |  | HIS 63 | 4.335015 |  |  |  |
|  |  |  |  | TYR82 | 4.81105 |  |  |  |
| **Shermilamine B**(Proposed Alkaloid) | -8.5 | His 63 | 5.17818 | PHE289 | 4.79301 | HIS 63 | | 4.98707 |
|  |  | His 63 | 4.51222 | - - | |  |  |  |
|  |  | His 63 | 4.07021 |  |  | HIS 63 | 4.12141 | |
|  |  | His 63 | 5.`14851 |  |  | ASP 48 | 3.59465 | |
|  |  | LEU64 | 5.31768 |  |  | GLU 241 | 3.50896 | |
|  |  |  |  |  |  | GLU 241 | 4.76763 | |
|  |  | PRO67 | 4.70543 |  |  |  |  |  |

**Table S5: ADMET properties of the standard and proposed drug**

| **Table S5: ADMET properties of the standard and proposed drug** | | | | |
| --- | --- | --- | --- | --- |
| **ADMET properties** | **Value for standard drug TLM** | **Probability for standard drug TLM** | **Value for Shermilamine B** | **Probability of Shermilamine B** |
| Human Intestinal Absorption | + | 0.9876 | + | 0.8744 |
| Caco-2 | + | 0.8584 | - | 0.8027 |
| Blood Brain Barrier | + | 0.9656 | + | 0.9828 |
| Human oral bioavailability | +  acceptable | 0.5143 acceptable | -  Not acceptable | 0.5000  Not acceptable |
| P-glycoprotein inhibitor | - | 0.9696 | + | 0.6496 |
| Carcinogenicity (binary) | -  acceptable | 0.6731  acceptable | -  acceptable | 0.8857  acceptable |
| Carcinogenicity (trinary) | Non-required  acceptable | 0.5295  acceptable | Non-required  acceptable | 0.6517  acceptable |
| Ames mutagenesis | -  acceptable | 0.7700  acceptable | +  Not acceptable | 0.6400 Not acceptable |
| Human either-a-go-go inhibition | - | 0.8139 | + | 0.7932 |
| CYP inhibitory promiscuity | - | 0.5616 | - | 0.6048 |
| Acute Oral Toxicity (c) | III | 0.7235 | III | 0.6424 |

Table S6: ADMET properties of the standard and proposed drug

| **Table S6: ADMET properties of the standard and proposed drug** | | | | |
| --- | --- | --- | --- | --- |
| **ADMET properties** | **Value for standard drug ZVT** | **Probability for standard drug ZVT** | **Value for Brachystamide B** | **Probability of Brachystamide B** |
| Human Intestinal Absorption | + | 0.9127 | + | 0.9771 |
| Caco-2 | - | 0.6657 | - | 0.6480 |
| Blood Brain Barrier | + | 0.9769 | + | 0.9811 |
| Human oral bioavailability | - Not acceptable | 0.5571  Not acceptable | - Not acceptable | 0.6286  Not acceptable |
| P-glycoprotein inhibitior | + | 0.7317 | + | 0.8162 |
| Carcinogenicity (binary) | - acceptable | 0.7908 acceptable | - acceptable | 0.9429 acceptable |
| Carcinogenicity (trinary) | Non-required  acceptable | 0.4752  acceptable | Non-required  acceptable | 0.5911  acceptable |
| Ames mutagenesis | -  acceptable | 0.6900  acceptable | -  acceptable | 0.8400  acceptable |
| Human either-a-go-go inhibition | +  Not acceptable | 0.7981  Not acceptable | +  Not acceptable | 0.7710  Not acceptable |
| CYP inhibitory promiscuity | + | 0.9093 | + | 0.7623 |
| Acute Oral Toxicity (c) | III | 0.5701 | 0.6716 | III |

Table S7: Physiochemical properties of the standard and proposed drugs

| **Table S7: Physiochemical properties of the standard and proposed drugs** | | | | |
| --- | --- | --- | --- | --- |
| **ADMET properties** | **Value for standard drug ZVT** | **Probability for standard drug ZVT** | **Value for Brachystamide B** | **Probability of Brachystamide B** |
| Human Intestinal Absorption | + | 0.9127 | + | 0.9771 |
| Caco-2 | - | 0.6657 | - | 0.6480 |
| Blood Brain Barrier | + | 0.9769 | + | 0.9811 |
| Human oral bioavailability | -  Not acceptable | 0.5571  Not acceptable | -  Not acceptable | 0.6286  Not acceptable |
| P-glycoprotein inhibitior | + | 0.7317 | + | 0.8162 |
| Carcinogenicity (binary) | -  acceptable | 0.7908  acceptable | -  acceptable | 0.9429  acceptable |
| Carcinogenicity (trinary) | Non-required  acceptable | 0.4752  acceptable | Non-required  acceptable | 0.5911  acceptable |
| Ames mutagenesis | -  acceptable | 0.6900  acceptable | -  acceptable | 0.8400  acceptable |
| Human either-a-go-go inhibition | +  Not acceptable | 0.7981  Not acceptable | +  Not acceptable | 0.7710  Not acceptable |
| CYP inhibitory promiscuity | + | 0.9093 | + | 0.7623 |
| Acute Oral Toxicity (c) | III | 0.5701 | 0.6716 | III |
